# Pivotal roles of PCNA loading and unloading on heterochromatin function

**DOI:** 10.1101/232181

**Authors:** Ryan Janke, Grant King, Martin Kupiec, Jasper Rine

## Abstract

In *Saccharomyces cerevisiae*, heterochromatin structures required for transcriptional silencing of the *HML* and *HMR* loci are duplicated in coordination with passing DNA replication forks. Despite major reorganization of chromatin structure, the heterochromatic, transcriptionally-silent states of *HML* and *HMR* are successfully maintained throughout S-phase. Mutations of specific components of the replisome diminish the capacity to maintain silencing of *HML* and *HMR* through replication. Similarly, mutations in histone chaperones involved in replication-coupled nucleosome assembly reduce gene silencing. Bridging these observations, we determined that the PCNA unloading activity of Elg1 was important for coordinating DNA replication forks with the process of replication-coupled nucleosome assembly to maintain silencing of *HML* and *HMR* through S-phase. Collectively these data identified a mechanism by which chromatin reassembly is coordinated with DNA replication to maintain silencing through S-phase.

**SIGNIFICANCE STATEMENT:** DNA replication poses a unique logistical challenge for the cell in that structural features of chromatin and their regulatory functions must be carefully coordinated with passage of replication machinery so faithful duplication of both the genome and its chromatin structures may be achieved. Nucleosome assembly is fundamental to reestablishment of chromatin in the wake of DNA replication, and here a mechanism by which nucleosome assembly is coordinated with DNA replication to maintain silenced chromatin is described.

## INTRODUCTION

Transcriptional repression within heterochromatin occurs through mechanisms that are typically insensitive to the identity of the genes encoded in the DNA of the repressed domains. The specific composition of proteins and epigenetic signatures that distinguish heterochromatin from euchromatin vary somewhat among species, and indeed even from one chromosome region to another. For example, in humans, regions of constitutive heterochromatin are typically enriched for H3K9 trimethylation whereas regions of facultative heterochromatin such as those found in inactivated X-chromosomes and at loci that regulate cell identity are typically enriched for H3K27 trimethylation (1, 2). Despite the range of mechanisms and proteins involved in heterochromatin formation and maintenance, certain characteristics appear universal and underlie a fundamental relationship between heterochromatin structure and function: heterochromatin is structurally compact, it localizes to distinct areas of the nucleus, and promotes transcriptional repression by limiting access of RNA Polymerases to DNA (3–10). The replication and epigenetic inheritance of heterochromatin is enigmatic in at least three ways. First, for heterochromatin that is epigenetically inherited, the unit of memory that allows for inheritance of the chromatin structure – and where it resides – is unclear. Second, the processes necessary to propagate that memory and reestablish the chromatin structure every cell cycle are poorly understood. Third, it is unclear how temporary and at least partial disassembly and reassembly of chromatin during DNA replication is coordinated and balanced with maintaining repression of genes in heterochromatin.

To gain further insight into how heterochromatin disassembly and reassembly during DNA replication occurs without loss of gene repression, we examined the well-characterized chromatin domains of the transcriptionally silent *HML* and *HMR* loci in *Saccharomyces cerevisiae*, which share structural features of heterochromatin in other eukaryotes. The chromatin at *HML* and *HMR* is composed of highly-ordered nucleosomes, each bound by the Sir2-Sir3-Sir4 complex, forming a compact heterochromatin structure necessary for transcriptional silencing (11). This structure is inherited, at least in part, through an epigenetic mechanism (12, 13). Establishment of silencing is initiated through the recruitment of the Sir complex to regulatory sites known as silencers that flank *HML* and *HMR* (14–16). Silencing is ultimately achieved upon Sir complex binding across *HML* and *HMR* where it deacetylates key positions on H3 and H4 N-terminal tails through the enzymatic activity of Sir2 (17–19). The resulting compact chromatin structure constrains access of RNA Pol II or Pol III at *HML* and *HMR* sufficiently to block transcription (10,20,21).

In dividing yeast cells, silencing is transiently lost in approximately 1 cell per 1000 cell divisions (22). This result implies that nearly all cells maintain silencing at *HML* and *HMR* through S-phase despite the need to replicate silent chromatin structure in the face of the partial nucleosome disassembly, and other chromatin changes, that occur during DNA replication. Hints at how the maintenance of silencing is balanced with DNA replication come from studies demonstrating that mutations in replication fork proteins and mutations affecting replication-coupled nucleosome assembly result in decreased silencing of *HML* and *HMR*. Replication-coupled nucleosome assembly is a multistep process that incorporates histones into newly replicated DNA within a few hundred bases from the replication fork (23, 24). Initially, Asf1 delivers newly synthesized H3-H4 dimers to replication forks where the H3-H4 dimers are transferred either to the CAF-1 complex or Rtt106, both of which deposit the histone dimers onto newly replicated DNA (25–29). Mutants of *asf1Δ*, *rtt106Δ*, and *cac1Δ* result in decreased silencing of *HML* and *HMR*, demonstrating the importance of proper chromatin assembly for heterochromatin function (30–38). CAF-1, a heterotrimeric histone chaperone composed of Cac1, Cac2, and Cac3 is coupled to DNA replication through a physical interaction with PCNA (39–41). PCNA travels with replication forks on both the leading and lagging strands where it increases the processivity of replicative DNA polymerases by physically tethering them to DNA, and also serves as an interaction scaffold to coordinate proteins involved in telomere maintenance, chromatin assembly, DNA-damage-response signaling and DNA repair with the replication fork (42). Certain mutations of *POL30*, which encodes PCNA, including some mutations that disrupt the physical interaction between PCNA and CAF-1, disrupt silencing at *HML*, *HMR*, and at telomeres (30, 39, 43). These data suggest that coordination of nucleosome assembly machinery with DNA replication forks is important to maintain silencing. PCNA is regulated, in part, by controlling when and where it resides on chromatin through the combined actions of two five-membered protein complexes that load and unload PCNA onto DNA (44). The RFC complex consisting of Rfc1-5, loads PCNA, and the other complex, Elg1-Rfc2-5, unloads PCNA (44–48). The PCNA unloading activity of Elg1 is important for multiple chromatin-based processes such as DNA repair, telomere-length maintenance, and telomeric silencing (49–51). *elg1Δ* mutants do not appear to impair Okazaki fragment processing or completion of DNA replication (47). Thus, the phenotypes caused by *elg1Δ* are not solely explained by DNA replication defects.

To drill into the processes that allow duplication of heterochromatin in particular, and all chromatin structures in general, we tested the extent to which proteins that act at the replication fork were necessary for the faithful propagation of heterochromatin’s gene silencing capability. Elg1 stood out as a major determinant of heterochromatin stability. This study revealed the mechanistic basis of Elg1’s contribution to heterochromatin stability, providing insight into the coordination between DNA replication and chromatin assembly by the factors that promote or limit PCNA.

## MATERIALS AND METHODS

### Strain and plasmids

All strains used in this study were derived from W303 (Table 1). CRASH assay strains, which use the switch from RFP to GFP expression upon derepression of *HML::cre*, were generated as described previously (22). Gene deletions were generated by integration of PCR-amplified disruption cassettes and confirmed by PCR using primers to amplify across the junctions at the site of integration (52, 53). The *pol30-D150E* point mutation was generated using Cas9 technology with the guide RNA sequence: 5′ aaacAATTCTCTAAAATTGTTCGTaa 3′, as described previously (54). The repair template was generated by annealing the oligo 5′ GTACGACTCCACCCTGTCATTGCCATCTTCTG<u>AATTCTCTAAAATTGTTCGTG</u> 3′ to oligo 5′ GATATTAATAGAATCACTCAATTGGGACAATT<u>CACGAACAATTTTAGAGAATT</u> 3′ and subsequently extending the 3′ ends using Phusion Polymerase (New England Biolabs, Ipswitch, MA).

**Table 1.**
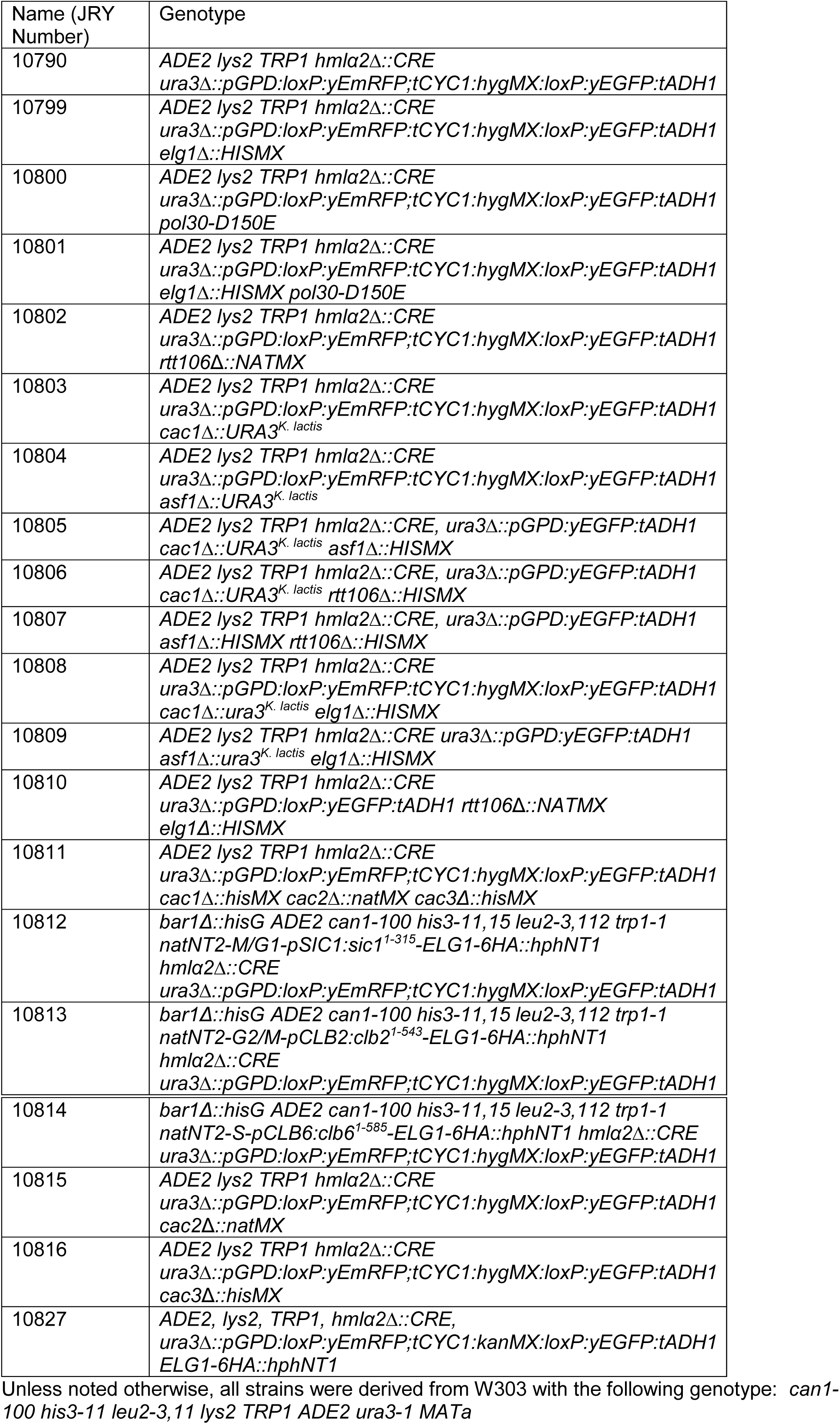
Strains used in this study

To generate plasmid pJR3418, the open reading frames and 5′ and 3′ UTRs of *CAC1*, *CAC2*, and *CAC3* were PCR amplified from genomic DNA using Gibson Assembly-compatible primers. The PCR fragments were assembled with plasmid pRS425 (55) linearized by SmaI digestion (New England Biolabs, Ipswitch, MA) using Gibson Assembly (New England Biolabs, Ipswitch, MA). To generate *ASF1* and *RTT106* overexpression plasmids, the open reading frame and 5′ and 3′ UTR of *RTT106* and *ASF1* were PCR amplified from genomic DNA using Gibson-Assembly compatible primers. The PCR fragments were assembled with pRS425 (55) linearized by SmaI digestion using Gibson Assembly to generate pJR3419 (*RTT106*) and pJR3425 (*ASF1*).

### Colony Growth and Imaging

Colonies were plated onto 1.5% agar plates containing yeast nitrogen base (YNB) without amino acids (Difco-Beckton-Dickinson, Franklin Lakes, NJ), 2% dextrose, and supplemented with complete supplement mixture (CSM)-Trp or CSM-Trp-Leu (Sunrise Science Products, San Diego, CA), as indicated. Colonies were incubated 5-7 days at 30°C. Colonies were imaged as described previously (56).

### Quantification of Silencing Loss by Flow Cytometry

For each CRASH strain, ten single colonies were inoculated separately into 2 ml of liquid yeast extract-peptone-2% dextrose (YPD) or CSM-Trp-Leu media in 96-deep-well plates and grown overnight to saturation at 30°C on a microplate orbital shaker. Overnight cultures were diluted into 1 ml of fresh media at a density of 10^5^ cells/ml in 96-deep-well plates and were grown at 30°C on a microplate orbital shaker until mid-log phase. For each culture, a minimum of 50,000 events were collected using an Attune NxT Flow Cytometer (Life Technologies). Scatterplots of forward scatter (height) and forward scatter (width) measurements were generated and gating was established to include only unbudded and budded cells and exclude debris and clumped cells for further analysis. Gating was used to measure separately the number of GFP-positive cells and the number of RFP-positive cells. Finally, a Boolean logic gate ‘RFP+ AND GFP+’ was used to determine the number of cells that were both GFP and RFP fluorescent. Such cells were inferred to have very recently undergone the Cre-mediated recombination event leading to GFP expression, yet retained RFP expressed in the recent past. The number of cells in a population that had very recently lost silencing (cells that were both GFP- and RFP-fluorescent) was divided by the number of cells in the population that had the potential to lose silencing (cells that were only RFP-fluorescent plus cells that are both GFP-and RFP-fluorescent) to obtain an apparent rate of silencing loss. Perdurance of RFP molecules in cells for 2-3 generations after switching from expression of RFP to GFP precluded direct calculation of true rates of silencing loss using this method. However, because the apparent rates of silencing loss directly reflected true silencing-loss rates, these values could be used to quantitatively compare silencing-loss rates across different genetic backgrounds. Boxplots were generated where the red line represents the median value calculated from at least 10 cultures. The blue boxes represent the 25^th^ and 75^th^ percentile. Whiskers represent the range of values within 1.5-times the interquartile range. Values extending beyond 1.5-times the interquartile range are marked as outliers (Red +). Unpaired two-sided (Student’s) *t* tests were used to determine whether differences in frequency of silencing loss were statistically significant.

### Chromatin Enrichment from Whole-Cell Extracts

An equivalent of 10 OD_600_ units (1 OD = ~10^7^ cells/ml) of cells were resuspended in 100 mM PIPES buffer (pH = 9.4) with 10 mM dithiothreitol (DTT) and incubated for 10 minutes at room temperature. The cells were pelleted by centrifugation and resuspended in 1 ml of spheroplast buffer containing 0.6 M sorbitol, 20 mM potassium phosphate (pH = 7.4), 5 mM DTT. To generate spheroplasts, zymolyase T100 was added at a final concentration of 80 μg/ml and the cells were incubated at 37°C for 10 minutes. The cells were repeatedly pelleted and washed once with 1 ml spheroplast buffer, and twice with 1 ml of a wash solution of 50 mM HEPES, 100 mM potassium chloride, 2.5 mM magnesium chloride, and 0.4 M sorbitol. The spheroplasts were resuspended in 200 μl of lysis buffer containing 50 mM HEPES (pH = 7.5), 100 mM potassium chloride, and 2.5 mM magnesium chloride, and protease inhibitor cocktail. To lyse spheroplasts, Triton X-100 was added to a final concentration of 0.25% and incubated for 5 minutes on ice with vortexing every minute. 50 μl of the whole-cell extract was gently pipetted on top of a 33% sucrose-lysate buffer solution and the chromatin fraction was separated by centrifugation at 12,000 RPM. The supernatant and chromatin pellet were collected separately and the chromatin was washed with lysis buffer. The fractions were mixed with Laemmli buffer and subject to separation by SDS-PAGE.

### Immuno-blotting

Samples were run on 4-20% gradient SDS-PAGE gels (Biorad, Hercules, CA) and transferred to nitrocellulose membranes (EMD-Millipore, Billerica, MA). The membranes were blocked in LI-CORE Odyssey Blocking Buffer (LI-CORE Biosciences, Lincoln, NE) and the following antibodies were used for immunodetection: anti-histone H3 (Ab1791, Abcam, Cambridge, MA), anti-PCNA (Ab221196, Abcam, Cambridge, MA), anti-phosphoglycerate kinase (no. 459250, Invitrogen, Camarillo, CA) and anti-FLAG (F3165, Sigma, St. Louis, MO). Membranes were incubated with infrared dye-conjugated antibodies IRDye800CW goat anti-mouse antibody and IRDye680RD goat anti-rabbit antibody (LI-CORE Biosciences, Lincoln, NE). Membranes were imaged using a LI-CORE Odyssey scanner in the 700 nm and 800 nm channels.

## RESULTS

### Elg1 was necessary for full silencing at *HML*

We measured silencing at *HML* using the previously described CRASH (Cre-reported altered states of heterochromatin) assay (22, 57). Briefly, the cre recombinase gene resides at the transcriptionally silent *HML* locus (Figure 1A). In these strains, even transient expression of Cre due to a loss of silencing at *HML* leads to site-specific recombination between LoxP sites and a permanent switch from RFP to GFP expression. Loss-of-silencing events, which manifest as GFP-expressing sectors in otherwise RFP-expressing yeast colonies, are detected at low levels in wild-type cells (Figure 1B) (22). *elg1Δ* mutations caused a substantial increase in silencing loss in *elg1Δ* strains compared to wild type, suggesting that control of PCNA unloading by Elg1 was important for silencing (Figure 1B). It was possible to quantify the frequency of silencing loss captured by the CRASH assay with flow cytometry. In CRASH strains, cells that had recently lost silencing contained both RFP and GFP for approximately three cell cycles. We utilized this property for quantification because the frequency of cells in a population that have recently lost silencing is proportional to the rate of silencing loss in that population (see Materials and Methods). For each sample, we calculated the ratio of cells in the population that had recently lost silencing to the number of cells in the population that had the potential to switch from RFP to GFP. We referred to this ratio as the ‘frequency of silencing loss’. The frequency of silencing loss in *elg1Δ* mutants was 8-fold greater than in wild type (Figure 1C).

**Figure 1.**
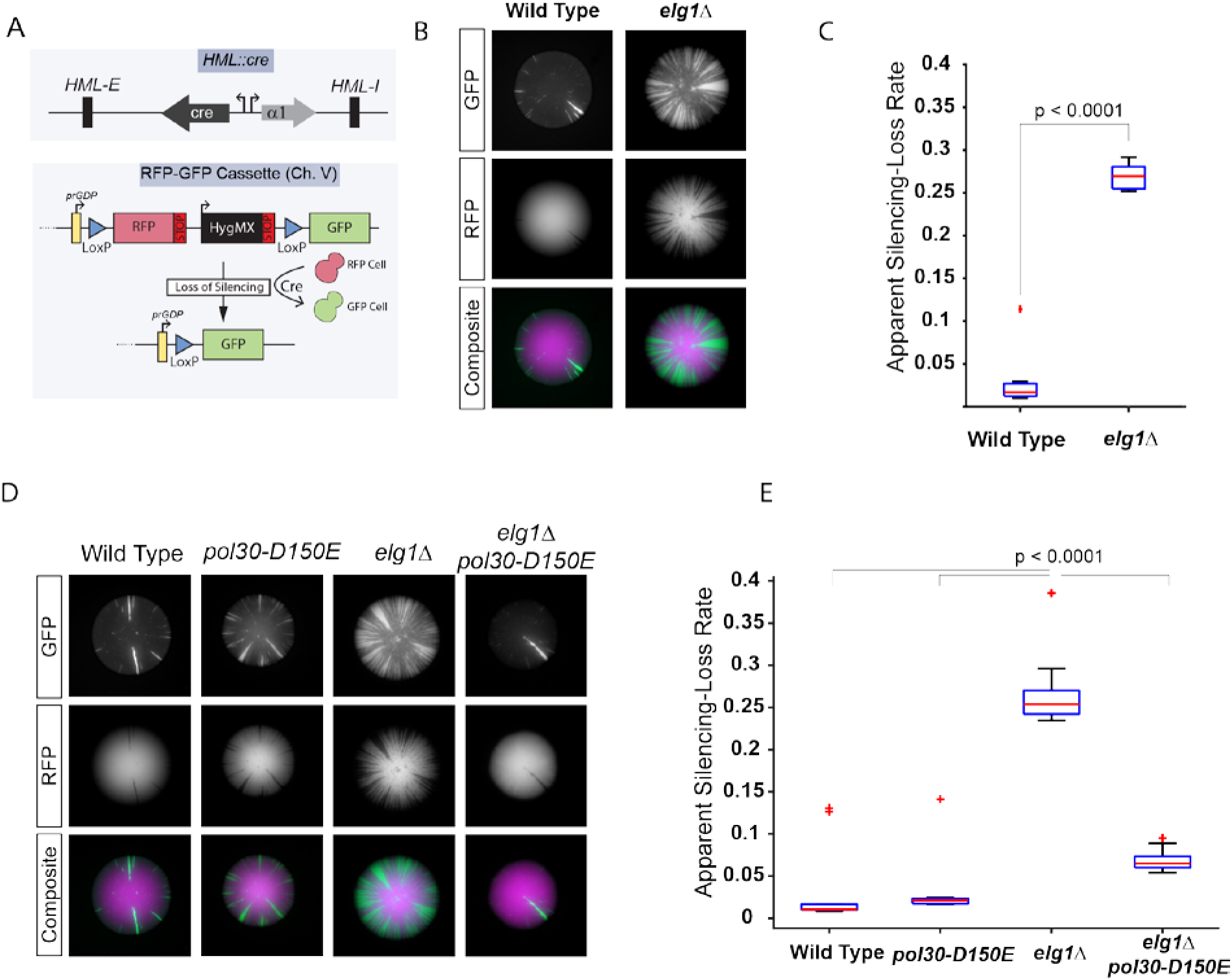
The PCNA unloading activity of Elg1 contributed to silencing *HML*. **A)** Illustration of the CRASH assay to measure silencing of *HML*. The CRASH assay contains two features: 1) Cre inserted at the *HML* locus controlled by the α2 promoter, 2) an RFP-GFP reporter cassette on chromosome V at the *URA3* locus. Cells that maintain silencing of *HML* constitutively express RFP driven by the *TDH3* promoter. Loss of silencing at *HML* leads to Cre expression, and results in Cre-mediated recombination between two LoxP sites resulting in removal of the RFP and HygMX genes, repositioning the GFP gene so that it becomes constitutively expressed. **B)** Representative images of colonies from wild-type (JRY10791) or *elg1Δ* mutant (JRY10799) strains containing the CRASH assay. **C)** The apparent silencing-loss rates of the strains in panel B were quantified by flow cytometry, as described in materials and methods. A *p*-value was calculated using an unpaired, two-tailed *t* test. **D)** Representative images of colonies from wild-type (JRY10791), *pol30-D150E* (JRY10800), *elg1Δ* (JRY10799) and *elg1Δ pol30-D150E* (JRY10801) mutant strains containing the CRASH assay. **E)** Quantification of apparent silencing-loss rates by flow cytometry of the strains in panel D. *p*-values were calculated by ANOVA and Tukey’s test post-hoc analysis.

It was formally possible that the increased loss of silencing associated with *elg1Δ* was due to an Elg1 function beyond its known role in unloading PCNA from DNA. To test whether or not the silencing defect was due to retention of PCNA on DNA, we utilized a previously characterized PCNA mutant (*pol30-D150E*) that disrupts the interaction interface between individual subunits of the PCNA trimer, causing it to spontaneously disassemble from chromatin even in the absence of Elg1 (58). These PCNA trimer-interface mutations rescue multiple other *elg1Δ* phenotypes (59). On its own, the *pol30-D150E* single mutant had no effect on silencing as measured by the CRASH assay, however the *pol30-D150E* mutant nearly completely rescued the silencing defect of an *elg1Δ* mutation (Figure 1D, 1E). These results implied that the silencing defects of *elg1Δ* were specifically due to the increased retention of PCNA on chromatin.

### Histone chaperones Asf1, CAF-1, and Rtt106 independently contributed to the maintenance of silencing at *HML*

How might increased retention of PCNA on chromatin in an *elg1Δ* mutant lead to a defect in silencing? Disruption of the interaction between CAF-1 and PCNA causes silencing defects, and PCNA acts as an important scaffold to control the recruitment of histone chaperone activity to the right time and place during S-phase (30, 39, 40). This led us to the hypothesis that removal of PCNA from chromatin following replication might be important for the regulation of histone chaperone function, and that failure to remove PCNA might impact nucleosome assembly and also silencing. Conventional genetic assays that measure the steady-state level of silencing at either *HML* or *HMR* averaged over a population of cells are capable of detecting silencing defects in *cac1Δ* mutants, but not in *asf1Δ* and *rtt106Δ* mutants. Instead, silencing phenotypes of *asf1Δ* and *rtt106Δ* mutants are detected only in genetic backgrounds that sensitize cells to silencing defects such as strains in which multiple histone chaperones are mutated (30, 41, 60, 61). These observations are consistent with two possible interpretations: *asf1Δ* and *rtt106Δ* mutants lack silencing phenotypes at *HML* and *HMR* because other histone chaperones compensate in those mutants; or the lack of phenotype reflects insufficient sensitivity of the assays. To distinguish between these two possibilities, the effect of mutant histone chaperones on silencing at *HML* was assessed with the CRASH assay, which is tuned to detect transient effects that reveal the dynamics of silenced chromatin (22). Deletion of *CAC1*, which encodes the largest subunit of the CAF-1 complex, resulted in increased loss of silencing (Figure 2A). Both *asf1Δ* and *rtt106Δ* mutations also resulted in increased silencing loss, but to a lesser degree than did CAF-complex mutants (Figure 2A, 2C). Thus individual histone chaperones independently contributed to silencing in ways that escaped detection by previous silencing assays.

**Figure 2.**
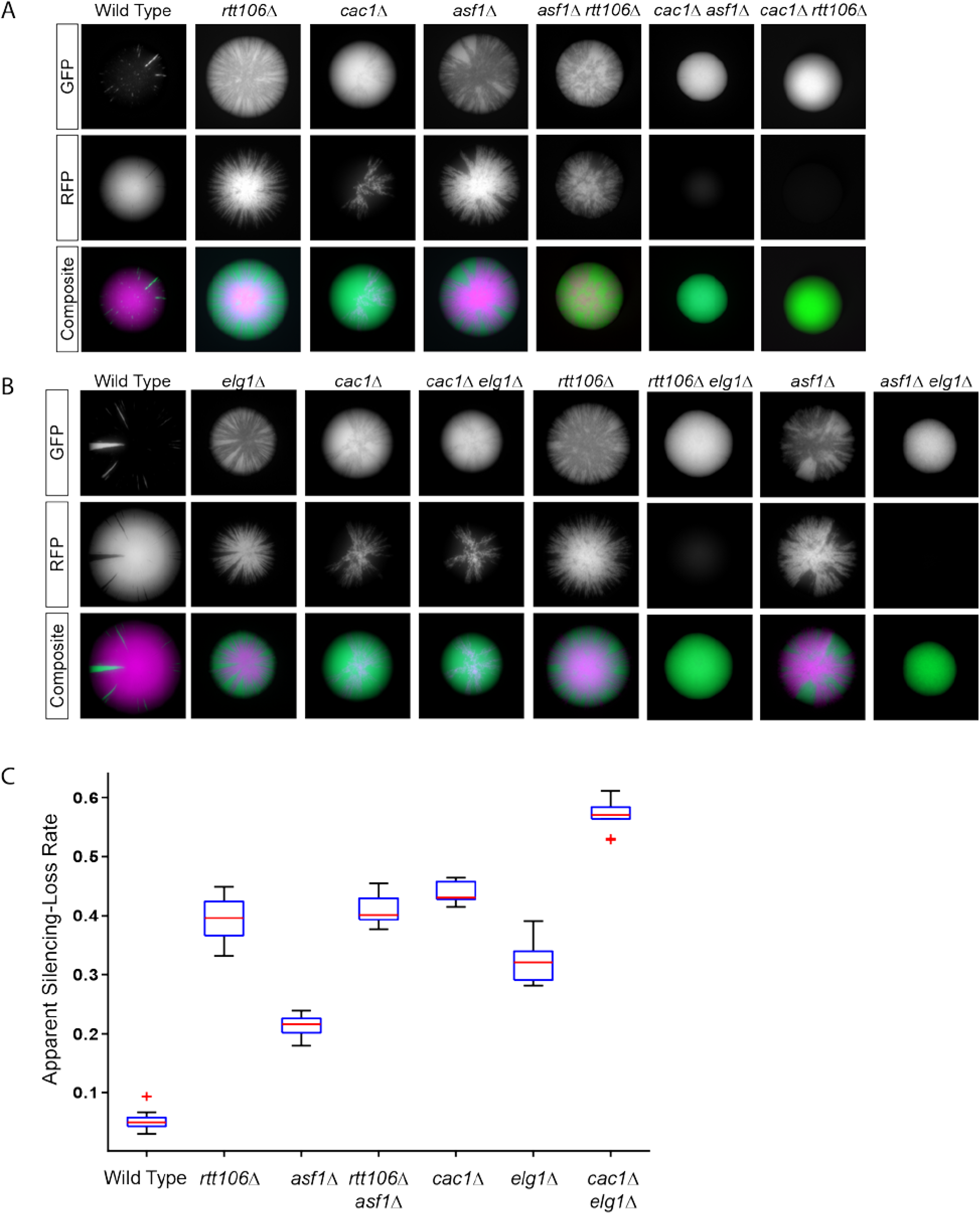
Genetic analysis of histone chaperone CAF-1 and Elg1 function in silencing. **A)** Representative images of colonies from wild type (JRY10791), *rtt106Δ* (JRY10802), *cac1Δ* (JRY10803), and *asf1Δ* (JRY10804) single mutants as well as representative images of colonies from the double mutants *cac1Δ asf1Δ* (JRY10805), *cac1Δ rtt106Δ* (JRY10806), and *asf1Δ rtt106Δ* (JRY10807), all of which carried the two components of the CRASH assay, are shown. Double-mutant strains of *cac1Δ rtt106Δ* and *cac1Δ asf1Δ* obtained from crosses of the single mutants uniformly expressed GFP, indicating they were unable to maintain silencing. **B)** Representative images of colonies with *elg1Δ* (JRY10799) in combination with the same histone chaperone single mutants as in panel A. *elg1Δ* (JRY10799) was crossed to histone chaperone mutants to obtain *elg1Δ cac1Δ* (JRY10808), *elg1Δ asf1Δ* (JRY10809), and *elg1Δ rtt106Δ* (JRY10810) double mutants. **C)** Quantification of apparent silencing-loss rates by flow cytometry of the single mutant strains, *elg1Δ cac1Δ, and asf1Δ rtt106Δ* double mutants from panel B. The rates of silencing loss of double mutant strains that uniformly expressed GFP could not be quantified.

Double-mutant analysis was performed to determine whether the histone chaperones function to maintain silencing through common or distinct pathways. In the case of *cac1Δ asf1Δ* and *cac1Δ rtt106Δ* double mutants, silencing defects were sufficiently strong that only GFP-expressing strains were recovered. These double mutants confer further increase in silencing loss compared to each of the single mutants (Figure 2A). Silencing defects in *asf1Δ rtt106Δ* double mutants were equivalent to *rtt106Δ* single mutants. These results were consistent with a model in which Asf1 and Rtt106 contribute to silencing through the same pathway. Therefore, all three histone chaperones contributed to the stability of silencing with the CAF-1 complex functioning in a distinct genetic pathway from Asf1 and Rtt106.

### Elg1 and the CAF-1 complex functioned in the same pathway to maintain silencing

The histone chaperone activity of the CAF-1 complex depends on its physical interaction with PCNA at replication forks (40, 43). The genome-wide distribution of PCNA bound to chromatin coincides with regions of active replication, which depends on a cycle of PCNA loading at primer-template junctions, including at the initiation of every Okazaki fragment, and rapid unloading from recently replicated regions (46, 62, 63). This pattern is disrupted in *elg1Δ* mutants in which PCNA is retained broadly across replicated regions of chromatin (46). We reasoned it was possible that proteins that interact with PCNA, such as CAF-1, might also become abnormally distributed across chromatin in the absence of Elg1. If the silencing phenotype of an *elg1Δ* mutant stems from inadequate CAF-1 function, the phenotype of a *cac1Δ* mutation on silencing would be qualitatively similar to that of *elg1Δ* (*i.e.* both mutations would cause silencing defects), and a *cac1Δ elg1Δ* double mutant would appear no more defective in silencing than a *cac1Δ* single mutant. In contrast if Elg1 and CAF-1 were to function in silencing through separate pathways, the silencing phenotype of a *cac1Δ elg1Δ* double mutant phenotype would be predicted to be substantially stronger than either of the single mutants. There was only a slightly higher increase (1.3-fold) in silencing loss in the *cac1Δ elg1Δ* double mutant compared to a *cac1Δ* single mutant (Figure 2B, C). In contrast, both *asf1Δ elg1Δ* and *rtt106Δ elg1Δ* double-mutant spores (derived from a cross of the single mutants) all lost silencing within the first few cell divisions (all double mutants, but not the single mutants or wild-type segregants, formed uniformly green colonies). Therefore, defects in silencing caused by *asf1Δ* and *rtt106Δ* were at least additive with the silencing defect of *elg1Δ*. We conclude that *CAC1* and *ELG1* function together to maintain silencing through a contribution to chromatin assembly that paralleled, but was distinct from, that of *ASF1* and *RTT106*. The slight increase in a *cac1Δ elg1Δ* double-mutant phenotype over that of the *cac1Δ* single mutant suggested that Elg1 also contributed to the maintenance of silencing through another process yet to be identified.

### CAF-1 and PCNA retention on chromatin increased in the absence of Elg1

In the absence of Elg1, the amount of chromatin-bound PCNA increases and PCNA remains on nascent DNA beyond S-phase (45, 59). Because CAF-1 physically interacts with PCNA, we tested whether the absence of Elg1 had a similar effect on CAF-1 association with chromatin. To measure binding of CAF-1 to chromatin, lysates were prepared from cycling cells and separated into chromatin and soluble fractions by centrifugation through a sucrose cushion. As expected, histone H3 was detected by immunoblot almost exclusively in the chromatin fraction (Figure 3A). Similar to previous observations, the amount of PCNA bound to chromatin increased by more than 50% in *elg1Δ* mutant cells compared to wild-type cells (Figure 3A, 3B). Importantly, the proportion of Cac1 retained in the chromatin fraction also increased significantly in *elg1Δ* mutant cells compared to wild-type cells (Figure 3A, 3B). No changes in the proportion of histone H3 bound to chromatin were observed in *elg1Δ* mutants relative to wild-type cells, which suggested that the enrichment of CAF-1 (and PCNA) in the chromatin fraction caused by *elg1Δ* mutants was not attributed to a generalized, non-specific increase of all chromatin binding proteins in those samples. Thus, in the absence of PCNA unloading, CAF-1 complex was retained on chromatin in excess of that in wild type.

**Figure 3.**
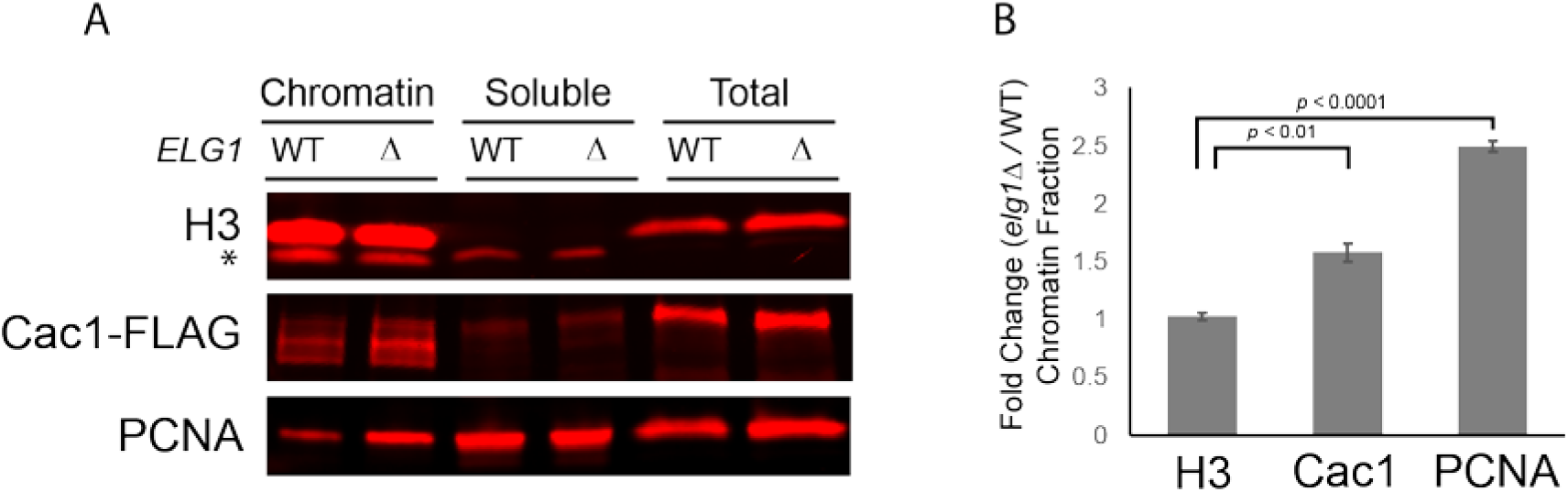
CAF-1 association with chromatin increased in an *elg1Δ* mutant. **A)** Immuno-blot analysis of histone H3, Caf1-FLAG, and PCNA levels in different subcellular fractions. Lysates prepared from wild-type (JRY10791) and *elg1Δ* (JRY10799) cells were separated into chromatin-associated material and soluble material. * indicates non-specific bands. **B)** Quantification of the immuno-blot signals of the chromatin fraction from panel A. The average fold-change values were calculated by dividing chromatin-associated levels measured in *elg1Δ* mutants by the levels measured in wild-type cells (n=3 biological replicates). Error bars represent standard error of the mean. p-values were calculated using unpaired, two-tailed *t* tests.

### Overexpression of the CAF-1 complex rescued *elg1Δ* silencing defect

In *elg1Δ* mutants, co-retention of CAF-1 with PCNA on DNA might cause a significant fraction of the CAF-1 pool to become sequestered from active replication forks where it functions. There are an estimated ~6,800 homotrimeric PCNA molecules and ~200-500 molecules of the CAF-1 complex in the nucleus (64). Because of a 13-30-fold excess of PCNA compared to CAF-1, CAF-1 may become limiting, leading to silencing defects observed in *elg1Δ*, yet enough free PCNA may be available at replication forks to support cell division. A simple prediction from this model is that overexpression of the CAF-1 complex should rescue silencing defects caused by *elg1Δ* mutants. To test this, a high-copy (2-micron) plasmid (pCAF-1) designed to overexpress all three proteins of the CAF-1 complex (Cac1-Cac2-Cac3) in a 1:1:1 stoichiometry was introduced into wild type and an *elg1Δ* mutant. No phenotype was observed in wildtype cells containing pCAF-1; excess CAF-1 complex did not interfere with normal silencing at *HML* (Figures 4C, 5A). As expected, pCAF-1 plasmid rescued the silencing defects of the *cac1Δ cac2Δ cac3Δ* triple mutant as well as *cac1Δ*, *cac2Δ*, and *cac3Δ* single mutants (Figures 4B, 4C, 5B) which demonstrated that pCAF-1 expressed functional CAF-1 complex. Importantly, pCAF-1 rescued the silencing defect of *elg1Δ* to nearly the same degree as it did with the *cac1Δ cac2Δ cac3Δ* triple mutant (Figures 4A, 4C). In contrast, overexpression of *ASF1* or *RTT106* was unable to restore silencing in *elg1Δ* mutants (Figures 4A, 4C) suggesting that the *elg1Δ* silencing defect was caused specifically by limited CAF-1 and not by limited histone chaperone activity per se. Overexpression of *ASF1* further weakened silencing in *elg1Δ* mutant (Figures 4A, 4C) and in wild-type cells (Figures 5A, 5F), consistent with previously reported effects of *ASF1* overexpression disrupting silencing at telomeres, *HML*, and *HMR* (36, 65).

**Figure 4.**
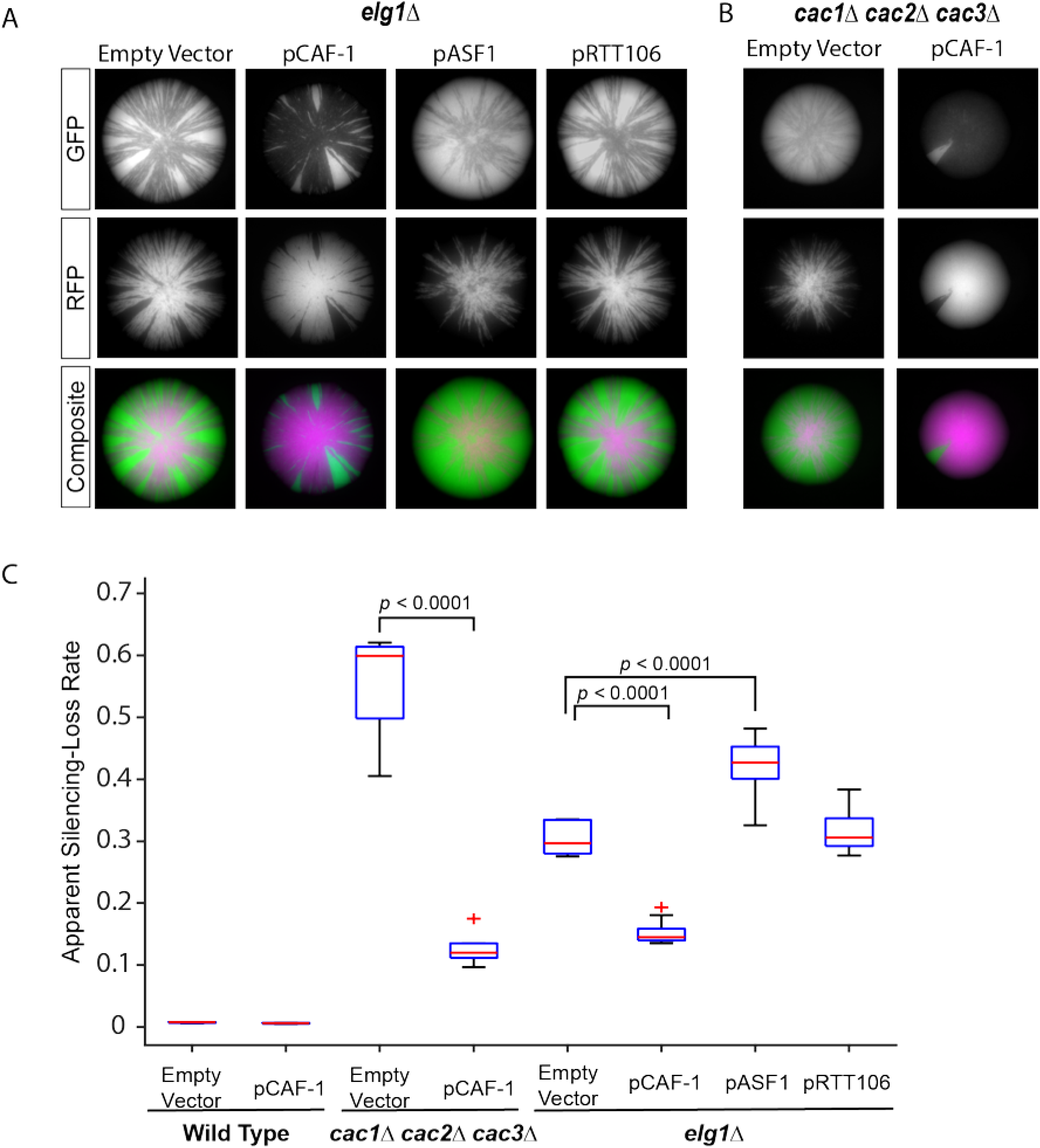
Overexpression of the CAF-1 complex rescued *elg1Δ* silencing defects. The empty-vector control pRS425 and a 2-micron high-copy plasmid containing either all three genes of the CAF-1 complex (pCAF-1, alias pJR3418), *ASF1* (pASF1, alias pJR3425), or *RTT106* (pRTT106, alias pJR3419) were transformed into CRASH assay strains. The cells were grown on a medium that maintained plasmid selection. Representative images of colonies containing the indicated plasmid in **A)** *elg1Δ* (JRY10799) and **B)** *cac1Δ cac2Δ cac3Δ* (JRY10811) strains. **C)** Plots of the apparent silencing-loss rates of wild type (JRY10791) transformed with pRS425 or pCAF-1 (pJR3418) and the strains in panels A and B quantified by flow cytometry. *p*-values were calculated by ANOVA and Tukey’s test post-hoc analysis.

**Figure 5.**
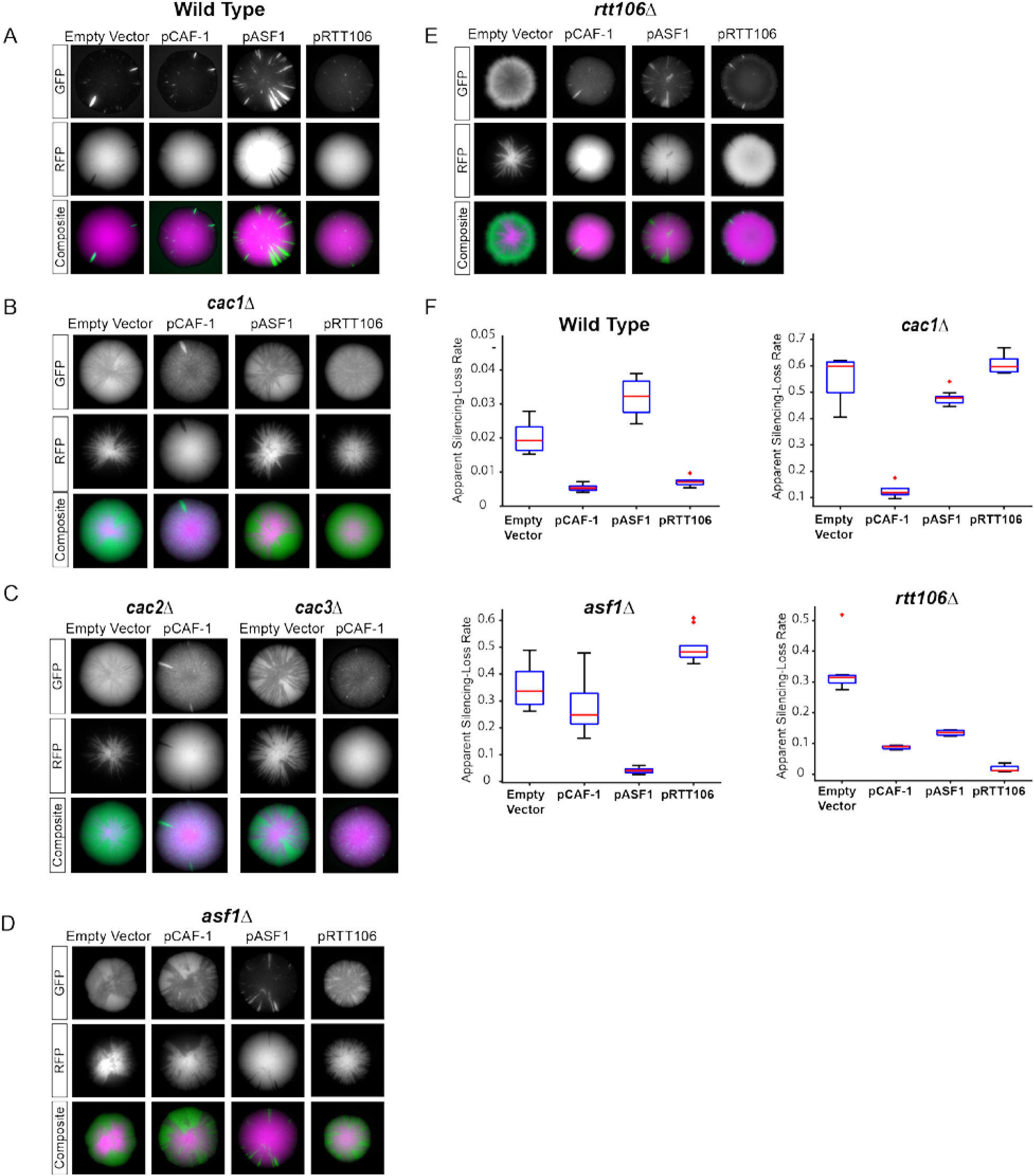
Impact of histone chaperone overexpression on silencing. **A)** Representative images of wildtype (JRY10791) colonies transformed with the empty-vector control pRS425 or a 2-micron high-copy plasmid containing either all three genes of the CAF-1 complex (pCAF-1, alias pJR3418), *ASF1* (pASF1, alias pJR3425), or *RTT106* (pRTT106, alias pJR3426). **B)** Images of *cac1Δ* (JRY10803) colonies transformed with the same plasmids as in A. **C)** Images of *cac2Δ* (JRY10815) and *cac3Δ* (JRY10816) colonies transformed with pRS425 or pCAF-1 (pJR3418). **D)** Images *asf1Δ* (JRY10804) colonies transformed with the same plasmids from panel A. **E)** Images of *rtt106Δ* (JRY10802) colonies transformed with the same plasmids from panel A. **F)** Plots of the apparent silencing-loss rates of the strains with the indicated mutations transformed with the plasmids described in panel A and quantified by flow cytometry.

### Quantitative and qualitative contributions of histone chaperones to gene silencing

The three histone chaperones tested in this study each contributed to the stability of silencing, with the CAF-1 complex functioning in a distinct genetic pathway from Asf1 and Rtt106. It is unknown how each histone chaperone contributes to maintaining silencing. If the silencing defects in individual histone chaperone mutants were caused by a general decrease in the overall capacity to deposit H3-H4 histones onto DNA, restoration of that biochemical activity, rather than the specific histone chaperone, would be expected to restore silencing. Alternatively, if deletion of a histone chaperone resulted in loss of a unique activity required for silencing specific to that protein, overexpression of other histone chaperones should not restore silencing. To distinguish between these models, silencing was measured in wild-type, *caf1Δ, asf1Δ*, and *rtt106Δ* strains transformed with high-copy (2-micron) histone chaperone expression plasmids. Overexpression of CAF-1 and *RTT106* had no effect on silencing in wild-type cells, while overexpression of *ASF1* destabilized silencing, as previously reported (Figures 5A, 5F) (36, 65). Overexpression of CAF-1, but not *RTT106* or *ASF1*, restored silencing in a *cac1Δ* mutant (Figures 4B, 5B, 5F). Similarly, silencing was restored in *asf1Δ* mutants by overexpression of *ASF1* but not CAF-1 or *RTT106* (Figures 5D, 5F). In *rtt106Δ* mutants, silencing was restored by overexpression by *RTT106* as well as CAF-1 and *ASF1* (Figures 5E,5F). Collectively these data were consistent with a model where Asf1 and CAF-1 contribute unique activities to silencing that cannot be supplanted by an excess of other histone chaperones, while the absence of Rtt106 could be compensated for by an overabundance of either Asf1 or CAF-1.

### Silencing at *HML* required S-phase-specific expression of Elg1

To determine if there was a specific cell-cycle stage in which Elg1-dependent unloading of PCNA was critical for silencing, the CRASH assay was introduced into strains that precisely control the expression and stability of Elg1 at specific cell-cycle stages (Figure 6A) (59). Expression of Elg1 during S-phase results in PCNA unloading activity while DNA replication takes place. In strains expressing Elg1 during G2/M or M/G1-phases, PCNA is retained on DNA from the last round of DNA replication until cells reach the cell-cycle stage where Elg1 is expressed. Silencing at *HML* was maintained to a much higher degree in cells when Elg1 was expressed in S-phase compared to cells in which Elg1 was expressed in M/G1-phase or G2/M-phase (Figure 6B). Although PCNA is unloaded when Elg1 is expressed during M/G1 or G2/M (59), this activity was insufficient to rescue the *elg1Δ* silencing defect. These results demonstrated that the PCNA unloading activity of Elg1 was necessary specifically during S-phase to ensure maximum maintenance of silencing at *HML*.

**Figure 6.**
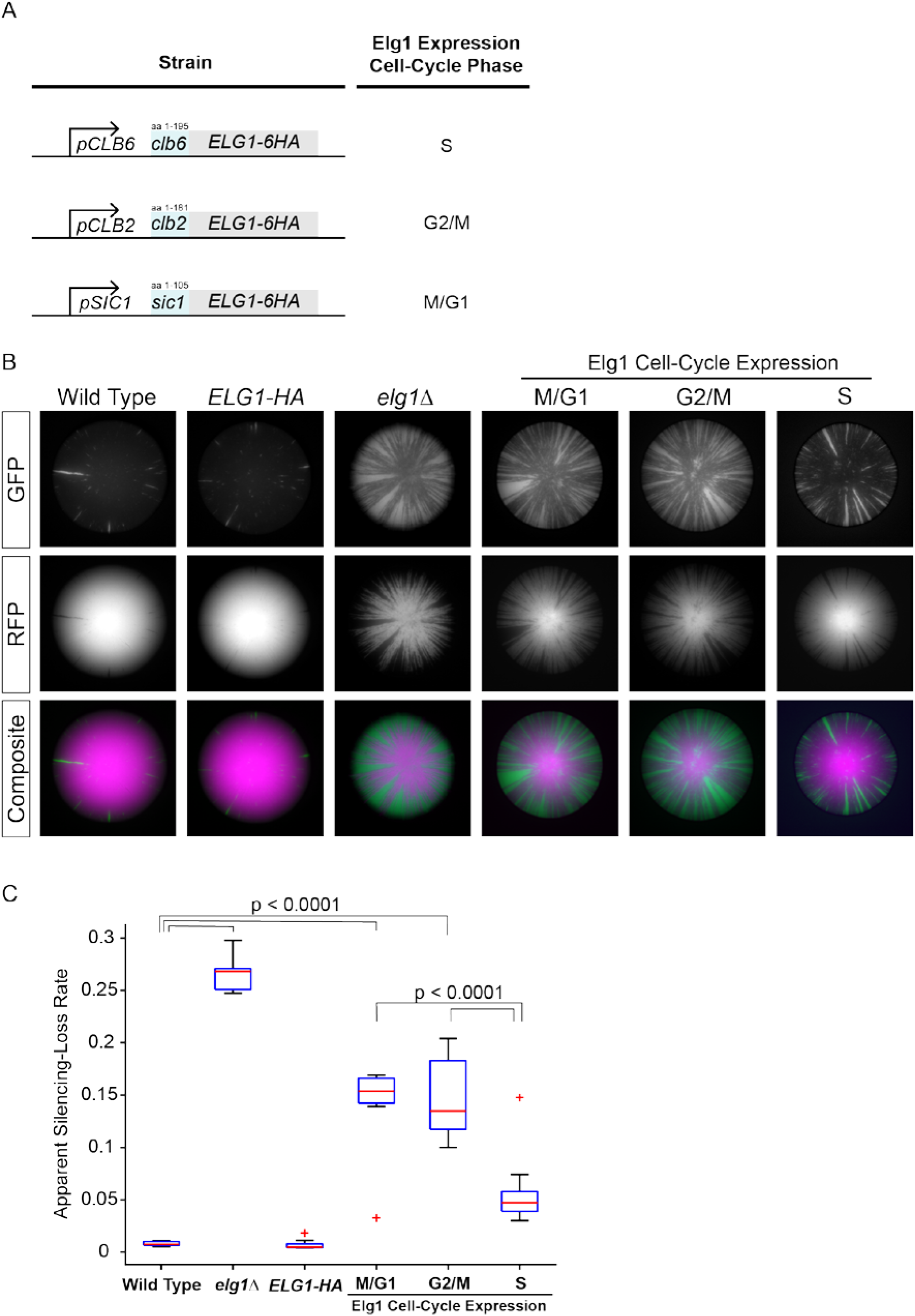
Cell-cycle-specific expression of Elg1 controlled the stability of gene silencing **A)** Diagram illustrating replacement of native *ELG1* promoter sequence and fusion of N-terminal degron sequences for cell-cycle-specific expression of Elg1. A Clb6-degron-Elg1 fusion protein expressed from a *CLB6* promoter is limited to S-phase (JRY10814), a Clb2-degron-Elg1 fusion protein expressed from a *CLB2* promoter is limited to G2/M-phase (JRY10813), and a Sic1-degron-Elg1 fusion expressed from a *SIC1* promoter is limed to M/G1-phase (JRY10812) (59). **B)** Representative colony images from wild type (JRY10791), an *ELG1-6HA* tagged strain (JRY10827), *elg1Δ* (JRY10799), or strains with cell-cycle-specific Elg1 expression as described in panel A. The cell-cycle expression stage of Elg1 is indicated. C) The apparent silencing-loss rates of the strains in panel B quantified by flow cytometry. *p*-values were calculated by ANOVA and Tukey’s test post-hoc analysis.

## DISCUSSION

This study revealed a previously uncharacterized role of Elg1 in S-phase for maintaining silenced chromatin and supported a model in which the PCNA unloading activity of Elg1 coordinates CAF-1-dependent nucleosome assembly with replication forks. PCNA serves as the scaffold that coordinates CAF-1 nucleosome assembly activity with DNA replication forks, and loss of the interaction between PCNA and CAF-1 disrupts CAF-1 nucleosome assembly function (30, 40, 43). Our data built upon this model and demonstrated that the function of CAF-1 also depends on maintaining PCNA’s ability to cycle on and off of DNA during replication. Several key observations supported such a model: First, double-mutant analysis demonstrated that silencing defects caused by loss of either *elg1Δ* or *cac1Δ* resulted from defects in a common process (CAF-1-dependent nucleosome assembly). Second, the silencing phenotype of an *elg1Δ* mutation could be suppressed in two ways: 1) through alleles that allow Elg1-independent unloading of PCNA by destabilizing the PCNA trimer ring (*pol30-D150E*) and 2) overexpression of the CAF-1 complex. These results genetically pinpointed persistence of PCNA on DNA as the cause for the silencing defect, and showed that this defect could be compensated for by more CAF-1 complex.

Molecular and cell-cycle experiments provided mechanistic support for this model. In the absence of Elg1, PCNA accumulates non-specifically on recently replicated chromatin in the wake of replication forks (46). We observed an increase in the fraction of PCNA bound to chromatin in *elg1Δ* mutants similar to the previously reported values. The fraction of CAF-1 associated with chromatin also increased significantly in the absence of Elg1. Controlled expression of Elg1 limited to S-phase was sufficient to maintain normal silencing, but Elg1 expression limited to G2/M or M/G1 phases resulted in defects in silencing that were similar to those of an *elg1Δ* mutant. A previous study that first reported the Elg1 promoter/degron cell-cycle-expression tools used here demonstrated that PCNA is efficiently unloaded from chromatin specifically during the intervals of Elg1 expression, but is retained on chromatin to excess when Elg1 is absent (59). Therefore, because silencing was maintained when Elg1 expression was limited to S-phase, the requirement for silencing to unload PCNA from chromatin was specific during S-phase. PCNA unloading during S-phase was critical for silencing presumably to ensure proper CAF-1 function, which acts specifically during replication-coupled chromatin assembly.

Why does PCNA need to cycle on and off DNA in S-phase? A combination of PCNA loading by Rfc1 and PCNA unloading by Elg1 ensures that PCNA is rapidly cycled on and off of chromatin during replication. PCNA is essential for viability because it is required to complete DNA replication, as is the activity of loading PCNA onto chromatin by Rfc1-5. In contrast, cells lacking Elg1 complete DNA replication with only a slight delay (10-15 minutes) compared to wild type (47, 66). Thus, Elg1 is not critical for the completion of DNA replication, per se. Instead, Elg1 may be critical for the cycling of PCNA on chromatin providing interaction sites for proteins that function specifically at replication forks and newly replicated DNA (40, 67). Unloading of PCNA in a timely manner by Elg1 may ensure that PCNA is present only at active sites of replication or recently replicated DNA where it coordinates the activity of proteins involved in DNA metabolism. In the absence of Elg1, widespread accumulation of PCNA on chromatin would preclude PCNA’s ability to serve as a marker that is specific for replicating DNA, resulting in miscoordination of PCNA-interacting proteins due to their recruitment to chromatin at the wrong time and place behind the replication fork.

We demonstrated that maintenance of the PCNA cycle by Elg1 is important for CAF-1 function. In the absence of Elg1, a larger fraction of CAF-1 became bound to chromatin. Because CAF-1 depends on interaction with PCNA for its recruitment to chromatin, it is possible that in *elg1* mutants, a significant fraction of CAF-1 becomes bound to PCNA that is not associated with active replication. There are estimated to be 200-500 CAF-1 molecules per cell and PCNA is estimated to be 13-30-fold more abundant than CAF-1 (64). However, considering there are approximately 350-400 annotated origins of replication in budding yeast, even if only a fraction of these fire during S-phase, a substantial portion of the CAF-1 pool (and potentially other PCNA-interacting proteins) may become sequestered with PCNA to chromatin to where the free pool of the CAF-1 complex becomes limiting at active replication forks, where it normally functions. Components of the CAF-1 complex are non-essential, and in this model, other histone chaperones would be adequate to support bulk chromatin assembly, but in the absence of CAF-1, the quality of that chromatin would be imperfect for heterochromatin formation and insufficient to maximally maintain silencing at *HML* and *HMR*. Consistent with this model, the silencing defects in an *elg1* mutant were suppressed by overexpression of the CAF-1 complex. Overall, our model suggests that one primary function of Elg1 is to maintain PCNA in a dynamic state during DNA replication so that PCNA and PCNA-bound proteins are able to cycle from nascent replicated DNA back to active sites of DNA replication.

Measurement of transient silencing loss revealed individual contributions of histone chaperones to silencing. The three histone chaperones that function during replication-coupled chromatin assembly, Asf1, Rtt106, and CAF-1, have known roles in silencing of *HML* and *HMR*. However, the individual contributions of each factor to silencing have been difficult to study, in part because of an overlap in function between the histone chaperones. Previous genetic studies of the contribution of histone chaperones towards silencing have mostly relied on transgene reporters that reflect steady state expression of either *HML* or *HMR* from a population of cells. Those reporters generally lacked sufficient sensitivity to detect effects of *asf1Δ* and *rtt106Δ* single mutants. The ability to measure transient losses of silencing with the CRASH assay uncovered the contribution of individual histone chaperones to silencing, and enabled double-mutant analysis that revealed which operated together and which operated at different steps. The inability to rescue the silencing phenotypes associated with loss of *ASF1* and *CAF-1* by overexpression of other histone chaperones established that each played a unique role in nucleosome assembly in heterochromatin that was not bypassed by increasing the abundance of other histone chaperones. Asf1 is required for H3K56 acetylation by Rtt109, a mark that stimulates the histone deposition activity of both CAF-1 and Rtt106 (27, 68). Both CAF-1 and Rtt106 deposit histone H3-H4 onto DNA. The silencing defect in an *rtt106Δ* mutant was rescued by overexpression of either *ASF1* or *CAF-1*, however the silencing defect in a *cac1Δ* mutant could not be rescued by *RTT106* overexpression. Thus CAF-1 fulfills a specialized role during nucleosome assembly, while the function of Rtt0106 could be met with the same biochemical activity of a different chaperone (in this case, CAF-1). An intriguing possibility is that CAF-1 has a specialized function in the assembly of nucleosomes on the lagging strand, where PCNA is more abundant.

DNA replication poses a unique logistical challenge for the cell in that structural features of chromatin and their regulatory functions must be carefully coordinated with passage of replication machinery so faithful duplication of both the genome and its chromatin structures may be achieved. Nucleosome assembly is fundamental to reestablishment of chromatin in the wake of DNA replication, and here we demonstrated that control of PCNA by Elg1 was necessary to coordinate nucleosome assembly to maintain transcriptional silencing through S-phase.

## ACKNOLWEDGEMENTS

We thank Takashi Kubota and Richard Kolodner for providing yeast strains and Paul Kaufman for sharing plasmids. This work was supported by NIH F32 fellowship (GM115074) to R.H.J., grants from the Berkeley-TAU fund to M.K. and J.R., from the Israel Science Foundation and the Volkswagen Foundation to M.K., NIH R01 grants (GM31105 and GM120374) to J.R.

